# TMbed – Transmembrane proteins predicted through Language Model embeddings

**DOI:** 10.1101/2022.06.12.495804

**Authors:** Michael Bernhofer, Burkhard Rost

**Author notes:** Corresponding author, https://www.rostlab.org/.

## Abstract

**Background:** Despite the immense importance of transmembrane proteins (TMP) for molecular biology and medicine, experimental 3D structures for TMPs remain about 4-5 times underrepresented compared to non-TMPs. Today’s top methods such as AlphaFold2 accurately predict 3D structures for many TMPs, but annotating transmembrane regions remains a limiting step for proteome-wide predictions.

**Results:** Here, we present TMbed, a novel method inputting embeddings from protein Language Models (pLMs, here ProtT5), to predict for each residue one of four classes: transmembrane helix (TMH), transmembrane strand (TMB), signal peptide, or other. TMbed completes predictions for entire proteomes within hours on a single consumer-grade desktop machine at performance levels similar or better than methods, which are using evolutionary information from multiple sequence alignments (MSAs) of protein families. On the per-protein level, TMbed correctly identified 94±8% of the beta barrel TMPs (53 of 57) and 98±1% of the alpha helical TMPs (557 of 571) in a non-redundant data set, at false positive rates well below 1% (erred on 30 of 5654 non-membrane proteins). On the per-segment level, TMbed correctly placed, on average, 9 of 10 transmembrane segments within five residues of the experimental observation. Our method can handle sequences of up to 4200 residues on standard graphics cards used in desktop PCs (e.g., NVIDIA GeForce RTX 3060).

**Conclusions:** Based on embeddings from pLMs and two novel filters (Gaussian and Viterbi), TMbed predicts alpha helical and beta barrel TMPs at least as accurately as any other method but at lower false positive rates. Given the few false positives and its outstanding speed, TMbed might be ideal to sieve through millions of 3D structures soon to be predicted, e.g., by AlphaFold2.

**Availability:** Our code, method, and data sets are freely available in the GitHub repository, https://github.com/BernhoferM/TMbed.

## Background

### Structural knowledge of TMPs 4-5 fold underrepresented

Transmembrane proteins (TMP) account for 20-30% of all proteins within any organism (1, 2); most TMPs cross the membrane with transmembrane helices (TMH). TMPs crossing with transmembrane beta strands (TMB), forming beta barrels, have been estimated to account for 1-2% of all proteins in Gram-negative bacteria; this variety is also present in mitochondria and chloroplasts (3). Membrane proteins facilitate many essential processes, including regulation, signaling, and transportation, rendering them targets for most known drugs (4, 5). Despite this immense relevance for molecular biology and medicine, only about 5% of all three-dimensional (3D) structures in the PDB (6, 7) constitute TMPs (8-10).

### Accurate 3D predictions available for proteomes need classification

The prediction of protein structure from sequence leaped in quality through AlphaFold2 (11), Nature’s method of the year 2021 (12). Although AlphaFold2 appears to provide accurate predictions for only very few novel “folds”, it importantly increases the width of structural coverage (13). AlphaFold2 seems to work well on TMPs (14), but for proteome-wide high-throughput studies, we still need to filter out membrane proteins from the structure predictions. Most state-of-the-art (SOTA) TMP prediction methods rely on evolutionary information in the form of multiple sequence alignments (MSA) to achieve their top performance. In our tests we included 13 such methods, namely BetAware-Deep (15), BOCTOPUS2 (16), CCTOP (17, 18), HMM-TM (19-21), OCTOPUS (22), Philius (23), PolyPhobius (24), PRED-TMBB2 (20, 21, 25), PROFtmb (3), SCAMPI2 (26), SPOCTOPUS (27), TMSEG (28), and TOPCONS2 (29).

### pLMs capture crucial information without MSAs

Mimicking recent advances of Language Models (LM) in natural language processing (NLP), protein Language Models (pLMs) learn to reconstruct masked parts of protein sequences based on the unmasked local and global information (30-37). Such pLMs, trained on billions of protein sequences, implicitly extract important information about protein structure and function, essentially capturing aspects of the “language of life” (32). These aspects can be extracted from the last layers of the deep learning networks into vectors, referred to as embeddings, and used as exclusive input to subsequent methods trained in supervised fashion to successfully predict aspects of protein structure and function (30-34, 36, 38-43). Often pLM-based methods outperform SOTA methods, which are using evolutionary information on top, and they usually require substantially fewer compute resources. Just before submitting this work, we became aware of another pLM-based TM-prediction method, namely DeepTMHMM (44) using ESM-1b (36) embeddings, and included it in our comparisons.

Here, we combined embeddings generated by the ProtT5 (34) pLM with a simple convolutional neural network (CNN) to create a fast and highly accurate prediction method for alpha helical and beta barrel transmembrane proteins and their overall inside/outside topology. Our new method, TMbed, predicted the presence and location of any TMBs, TMHs, and signal peptides for all proteins of the human proteome within 46 minutes on our server machine (SOM: Table S1) at the same or better level of performance as other methods, which require substantially more time.

## Materials & Methods

### Data set: membrane proteins (TMPs)

We collected all primary structure files for alpha helical and beta barrel transmembrane proteins (TMP) from OPM (45) and mapped their PDB (6, 7) chain identifiers (PDB-id) to UniProtKB (46) through SIFTS (47, 48). Toward this end, we discarded all chimeric chains, all models, and all chains for which OPM failed to map any transmembrane start or end position. This resulted in 2,053 and 206 sequence-unique PDB chains for alpha helical and beta barrel TMPs, respectively.

We used the ATOM coordinates inside the OPM files to assign the inside/outside orientation of sequence segments not within the membrane. We manually inspected inconsistent annotations (e.g., if both ends of a transmembrane segment had the same inside/outside orientation) and cross-referenced them with PDBTM (49-51), PDB, and UniProtKB. We then either corrected such inconsistent annotations or discarded the whole sequence. As OPM does not include signal peptide annotations, we compared our TMP data sets to the set used by SignalP 6.0 (52) and all sequences in UniProtKB/Swiss-Prot with experimentally annotated signal peptides using CD-HIT (53, 54). For any matches with at least 95% global sequence identity (PIDE), we transferred the signal peptide annotation onto our TMPs. We removed all sequences with fewer than 50 residues to avoid noise from incorrect sequencing fragments, and all sequences with over 15,000 residues to save energy (lower computational costs).

Finally, we removed redundant sequences from the two TMP data sets by clustering them with MMseqs2 (55) to at most 20% local pairwise sequence identity (PIDE) with 40% minimum alignment coverage, i.e., no pair had more than 20% PIDE for any local alignment covering at least 40% of the shorter sequence. The final non-redundant TMP data sets contained 593 alpha helical TMPs and 65 beta barrel TMPs, respectively.

### Data set: globular non-membrane proteins

We used the SignalP 6.0 (SP6) dataset for our globular proteins. As the SP6 dataset contained only the first 70 residues of each protein, we took the full sequences from UniProtKB/Swiss-Prot and transferred the signal peptide annotations. To remove any potential membrane proteins from this non-TMP data set, we compared it with CD-HIT (53, 54) against three other data sets: (1) our TMP data sets before redundancy reduction, (2) all protein sequences from UniProtKB/Swiss-Prot with any annotations of transmembrane segments, and (3) all proteins from UniProtKB/Swiss-Prot with any subcellular location annotations for membrane. We removed all proteins from our non-TMP data set with more than 60% global PIDE to any protein in sets 1-3. Again, we dropped all sequences with less than 50 or more than 15,000 residues and applied the same redundancy reduction as before (20% PIDE at 40% alignment coverage). The final non-redundant data set contained 5,859 globular, water-soluble non-TMP proteins; 698 of these have a signal peptide.

### Additional redundancy reduction

One anonymous reviewer spotted homologs in our data set after the application of the above protocol. To address this problem, we performed another iteration of redundancy reduction for each of the three data sets using CD-HIT at 20% PIDE. In order to save energy (i.e., avoid retraining our model), we decided to remove clashes for the evaluation, i.e., if two proteins shared more than 20% PIDE, we removed both from the data set (as TMbed was trained on both in the cross-validation protocol). Thereby, this second iteration removed 235 proteins: 8 beta barrel TMPs, 22 alpha helical TMPs, and 205 globular, non-membrane proteins. Our final test data sets included 57 beta barrel TMPs, 571 alpha helical TMPs, and 5,654 globular, non-membrane proteins.

### Membrane re-entrant regions

Besides transmembrane segments that cross the entire membrane, there are also others, namely membrane segments that briefly enter and exit the membrane on the same side. These are referred to as re-entrant regions (56, 57). Although rare, some methods explicitly predict them (17, 18, 22, 27, 58). However, as OPM does not explicitly annotate such regions and since our data set already had a substantial class imbalance between beta barrel TMPs, alpha helical TMPs and, globular proteins, we decided not to predict re-entrant regions.

### Embeddings

We generated embeddings with protein Language Models (pLMs) for our data sets using a transformer-based pLM ProtT5-XL-U50 (short: ProtT5) (34). We discarded the decoder part of ProtT5, keeping only the encoder for increased efficiency (note: encoder embeddings are more informative (34)). The encoder model converts a protein sequence into an embedding matrix that represents each residue in the protein, i.e., each position in the sequence, by a 1024-dimensional vector containing global and local contextualized information. We converted the ProtT5 encoder from 32-bit to 16-bit floating-point format to reduce the memory footprint on the GPU. We took the pre-trained ProtT5 model as is without any further task-specific fine-tuning.

We chose ProtT5 over other embedding models, such as ESM-1b (36), based on our experience with the model and comparisons during previous projects (34, 38). Furthermore, ProtT5 does not require splitting long sequences, which might remove valuable global context information, while ESM-1b can only handle sequences of up to 1022 residues.

### Model architecture

Our TMbed model architecture contained three modules (SOM: Fig. S1): a <u>convolutional neural network (CNN</u>) to generate per-residue predictions, a <u>Gaussian smoothing filter</u>, and a <u>Viterbi decoder</u> to find the best class label for each residue. We implemented the model in PyTorch (59).

#### Module 1: CNN

The first component of TMbed is a CNN with four layers (SOM: Fig. S1). The first layer is a pointwise convolution, i.e., a convolution with kernel size of 1, which reduces the ProtT5 embeddings for each residue (position in the sequence) from 1024 to 64 dimensions. Next, the model applies layer normalization (60) along the sequence and feature dimensions, followed by a ReLU (Rectified Linear Unit) activation function to introduce non-linearity. The second and third layers consist of two parallel depthwise convolutions; both process the output of the first layer. As depthwise convolutions process each input dimension (feature) independently while considering consecutive residues, those two layers effectively generate sliding weighted sums for each dimension. The kernel sizes of the second and third layer are 9 and 21, respectively, corresponding to the average length of transmembrane beta strands and helices. As before, the model normalizes the output of both layers and applies the ReLU function. It then concatenates the output of all three layers, constructing a 192-dimensional feature vector for each residue (position in the sequence). The fourth layer is a pointwise convolution combining the outputs from the previous three layers and generates scores for each of the five classes: transmembrane beta strand (B), transmembrane helix (H), signal peptide (S), non-membrane inside (i), and non-membrane outside (o).

#### Module 2: Gaussian filter

This module smooths the output from the CNN for adjacent residues (sequence positions) to reduce noisy predictions. The filter allows flattening isolated single-residue peaks. For instance, peaks extending of only one to three residues for the classes B and H are often non-informative; similarly short peaks for class S are unlikely correct. The filter uses a Gaussian distribution with standard deviation of 1 and a kernel size of 7, i.e., its seven weights correspond to three standard deviation intervals to the left and right, as well as the central peak. A softmax function then converts the filtered class scores to a class probability distribution.

#### Module 3: Viterbi decoder

The Viterbi algorithm decodes the class probabilities and assigns a class label to each residue (position in the sequence; SOM: Note S3, Fig. S2). The algorithm uses no trainable parameter; it scores transitions according to the predicted class probabilities. Its purpose is to enforce a simple grammar such that (1) signal peptides can only start at the N-terminus (first residue in protein), (2) signal peptides and transmembrane segments must be at least five residues long (a reasonable trade-off between filtering out false positives and still capturing weak signals), and (3) the prediction for the inside/outside orientation has to change after each transmembrane segment (to simulate crossing through the membrane). Unlike the Gaussian filter, we did not apply the Viterbi decoder during training. This simplified backpropagation and sped up training.

### Training details

We performed a stratified five-fold nested cross-validation for model development (SOM: Fig. S3). First, we separated our protein sequences into four groups: beta barrel TMPs, alpha helical TMPs with only a single helix, those with multiple helices, and non-membrane proteins. We further subdivided each group into proteins with and without signal peptides. Next, we randomly and evenly distributed all eight groups into five data sets. As all of our data sets were redundancy reduced, no two splits contained similar protein sequences for any of the classes. However, similarities between proteins of two different classes were allowed, not the least to provide more conservative performance estimates.

During development, we used four of the five splits to create the model and the fifth for testing (SOM: Fig. S3). Of the first four splits, we used three to train the model and the fourth for validation (optimize hyperparameters). We repeated this 3-1 split three more times, each time using a different split for the validation set, and calculated the average performance for every hyperparameter configuration. Next, we trained a model with the best configuration on all four development splits and estimated its final performance on the independent test split. We performed this whole process a total of five times, each time using a different of the five splits as test data and the remaining four for the development data. This resulted in five final models; each trained, optimized, and tested on independent data sets.

We applied weight decay to all trained weights of the model and added a dropout layer right before the fourth convolutional layer, i.e., the output layer of the CNN. For every training sample (protein sequence), the dropout layer randomly sets 50% of the features to zero across the entire sequence, preventing the model from relying on only a specific subset of features for the prediction.

We trained all models for 15 epochs using the AdamW (61) optimizer and cross-entropy loss. We set the beta parameters to 0.9 and 0.999, used a batch size of 16 sequences, and applied exponential learning rate decay by multiplying the learning rate with a factor of 0.8 every epoch. The initial learning rate and weight decay values were part of the hyperparameters optimized during cross-validation (SOM: Table S2).

The final TMbed model constitutes an ensemble over the five models obtained from the five outer cross-validation iterations (SOM: Fig. S3), i.e., one for each training/test set combination. During runtime, each model generates its own class probabilities (CNN, plus Gaussian filter), which are then averaged and processed by the Viterbi decoder to generate the class labels.

### Evaluation and other methods

We evaluated the test performance of TMbed on a per-protein level and on a per-segment level (SOM: Note S1). For protein level statistics, we calculated recall and false positive rate (FPR). We computed those statistics for three protein classes: alpha helical TMPs, beta barrel TMPs, and globular proteins.

We distinguished correct and incorrect segment predictions using two constraints: 1) the observed and predicted segment must overlap such that the intersection of the two is at least half of their union, and 2) neither the start nor the end positions may deviate by more than five residues between the observed and predicted segment (SOM: Fig. S4). All segments predicted meeting both these criteria were considered as “correctly predicted segments”, all others as “incorrectly predicted segments”. This allowed for a reasonable margin of error regarding the position of a predicted segment, while punishing any gaps introduced into a segment. For per-segment statistics, we calculated recall and precision. We also computed the percentage of proteins with the correct number of predicted segments (Q_num_), the percentage of proteins for which all segments are correctly predicted (Q_ok_), and the percentage of correctly predicted segments that also have the correct orientation within the membrane (Q_top_). We considered only proteins that actually contain the corresponding type of segment when calculating per-segment statistics, e.g., only beta barrel TMPs for transmembrane beta strand segments. We compared TMbed to other prediction methods for alpha helical and beta barrel TMPs (details in SOM Note S2): BetAware-Deep (15), BOCTOPUS2 (16), CCTOP (17, 18), DeepTMHMM (44), HMM-TM (19-21), OCTOPUS (22), Philius (23), PolyPhobius (24), PRED-TMBB2 (20, 21, 25), PROFtmb (3), SCAMPI2 (26), SPOCTOPUS (27), TMSEG (28), and TOPCONS2 (29). We chose those methods based on their good prediction accuracy and public popularity. For methods predicting only either alpha helical or beta barrel TMPs, we considered the corresponding other type of TMPs as globular proteins for the per-protein statistics. In addition, we generated signal peptide predictions with SignalP 6.0 (52). The performance of older TMH prediction methods could be triangulated based on previous comprehensive estimate of such methods (28, 62).

Unless stated otherwise, all reported performance values constitute the average performance over the five independent test sets during cross-validation (c.f. *Training details*) and their error margins reflect the 95% confidence interval (CI), i.e., 1.96 times the sample standard error over those five splits (SOM: Tables S5 & S6). We considered two values *A* and *B* statistically significantly different if they differ by more than their composite 95% confidence interval:

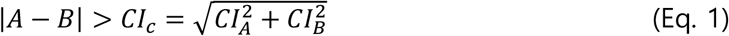

### Additional out-of-distribution benchmark

In the most general sense, machine learning models learn and predict distributions. Most membrane data sets are small and created using the same resources, including OPM (45), PDBTM (49-51), and UniProtKB/Swiss-Prot (46) that often mix experimental annotations with sophisticated algorithms (50, 63-65) to determine the boundaries of transmembrane segments, e.g., by using the 3D structure. Given these constraints, we might expect data sets from different groups to render similar results. Analyzing the validity of this assumption, we included the data set assembled for the development of DeepTMHMM (44). Three reasons made us chose this set as an alternative perspective: (1) it is recent, (2) it contains helical and beta barrel TMPs, and (3) the authors made their cross-validation predictions available, simplifying comparisons.

We created two distinct data sets from the DeepTMHMM data. First, we collected all proteins common to both data sets (TMbed and DeepTMHMM). We used those proteins to estimate how much the annotations within both data sets agree with each other. In total, there were 1,788 proteins common to both data sets: 43 beta barrel TMPs, 184 alpha helical TMPs, 1,560 globular proteins, and one protein (MSPA_MYCS2; Porin MspA) which sits in the outer-membrane of *Mycobacgterium smegmatis (66)*. We classified this as beta barrel TMP while DeepTMHMM listed it, most likely incorrectly, as a globular protein. The second data set that we created contained all proteins from the DeepTMHMM data set that were non-redundant to the training data of TMbed. We used PSI-BLAST (67) to find all significant (e-value < 10^−4^) local alignments with a 20% PIDE threshold and 40% alignment coverage to remove the redundant sequences. This second data set contained 667 proteins: 14 beta barrel TMPs, 86 alpha helical TMPs, and 567 globular proteins. We generated predictions with TMbed for those proteins and compared them to the cross-validation predictions for DeepTMHMM, as well as the best performing methods from our own benchmark (CCTOP (17, 18), TOPCONS2 (29), BOCTOPUS2 (16)); we used the DeepTMHMM data set annotations as ground truth.

### Data set of new membrane proteins

In order to perform a CASP-like performance evaluation, we gathered all PDB structures published since Feb 05, 2022, which is just after the data for our set and that of DeepTMHMM (44) have been collected. This comprised 1,511 PDB structures (more than 250 of which related to the SARS-CoV-2 protein P0DTD1) that we could map to 1,078 different UniProtKB sequences. We then used PSI-BLAST to remove all sequences similar to our data set or that of DeepTMHMM (e-value < 10^−4^, 20% PIDE at 40% coverage), which resulted in 333 proteins. Next, we predicted transmembrane segments within those proteins using TMbed and DeepTMHMM. For 38 proteins, either TMbed or DeepTMHMM predicted transmembrane segments. After removing any sequences shorter than 100 residues (i.e., fragments) and those in which the predicted segments were not within the resolved regions of the PDB structure, we were left with a set of 5 proteins: one beta barrel TMP and four alpha helical TMPs. Finally, we used the PPM (63-65) algorithm from OPM (45) to estimate the actual membrane boundaries.

## Results & Discussion

We have developed a new machine learning model, dubbed TMbed; it exclusively uses embeddings from the ProtT5 (34) pLM as input to predict for each residue in a protein sequence to which of the following four “classes” it belongs: transmembrane beta strand (TMB), transmembrane helix (TMH), signal peptide (SP), or non-transmembrane segment. It also predicts the inside/outside orientation of TMBs and TMHs within the membrane, indicating which parts of a protein are inside or outside a cell or compartment. Although the prediction of signal peptides was primarily integrated to improve TMH predictions by preventing the confusion of TMHs with SPs and *vice versa*, we also evaluated and compared the performance for SP prediction of TMbed to that of other methods.

### Reaching SOTA in protein sorting

TMbed detected TMPs with TMHs and TMBs at levels similar or numerically above the best state-of-the-art (SOTA) methods that use evolutionary information from multiple sequence alignments (MSA; Table 1: Recall). Compared to MSA-based methods, TMbed achieved this parity or improvement at a significantly lower false positive rate (FPR), tied only with DeepTMHMM (44), another embedding-based method (Table 1: FPR). Given those numbers, we expect TMbed to misclassify only about 215 proteins for a proteome with 20,000 proteins (SOM: Table S10), e.g., the human proteome, while the other methods would make hundreds more mistakes (DeepTMHMM: 331, TOPCONS2: 683, BOCTOPUS2: 880). Such low FPRs suggest our method as an automated high-throughput filter for TMP detection, e.g., for the creation and annotation of databases, or the decision which AlphaFold2 (11, 68) predictions to parse through advanced software annotating transmembrane regions in 3D structures or predictions (45, 49, 69). In the binary prediction of whether or not a protein has a signal peptide, TMbed achieved similar levels as the specialist SignalP 6.0 (52) and as DeepTMHMM (44), reaching 99% recall at 0.1% FPR (SOM: Table S3).

**Table 1:** Per-protein performance. *.

| | β-TMP (57) | | α-TMP (571) | | Globular (5,654) | |
| --- | --- | --- | --- | --- | --- | --- |
|  | Recall (%) | FPR (%) | Recall (%) | FPR (%) | Recall (%) | FPR (%) |
| TMbed | **93.8± 7.5** | **0.1±0.1** | **97.5±0.7** | **0.5±0.2** | **99.5±0.2** | 2.8± 1.2 |
| DeepTMHMM | 77.9±12.7 | **0.1±0.1** | 95.8±1.3 | **0.5±0.2** | **99.5±0.2** | 5.9± 2.2 |
| TMSEG | - | - | 96.5±1.0 | 2.3±0.3 | 97.7±0.3 | 3.5± 1.0 |
| TOPCONS2<sup>1</sup> | - | - | 94.2±1.3 | 2.6±0.3 | 97.4±0.3 | 5.8± 1.3 |
| OCTOPUS<sup>1</sup> | - | - | 94.2±1.9 | 9.1±0.7 | 90.9±0.7 | 5.8± 1.9 |
| Philius<sup>1</sup> | - | - | 92.5±1.4 | 2.6±0.2 | 97.4±0.2 | 7.5± 1.4 |
| PolyPhobius<sup>1</sup> | - | - | 97.2±1.1 | 5.3±0.4 | 94.7±0.4 | 2.8± 1.1 |
| SPOCTOPUS<sup>1</sup> | - | - | **97.5±1.6** | 17.2±0.8 | 82.8±0.8 | **2.5± 1.6** |
| SCAMPI2 (MSA) | - | - | 94.2±1.6 | 5.6±0.3 | 94.4±0.3 | 5.8± 1.6 |
| CCTOP<sup>2</sup> | | | 96.1±2.1 | 3.7±0.6 | 96.3±0.6 | 3.9± 2.1 |
| HMM-TM (MSA)<sup>3</sup> | - | - | 97.3±1.6 | 21.4±0.5 | 78.6±0.5 | 2.7± 1.6 |
| BOCTOPUS2 | 84.0±13.3 | 4.2±0.5 | - | - | 95.8±0.5 | 16.0±13.3 |
| BetAware-Deep | 85.1± 9.3 | 4.7±0.3 | - | - | 95.3±0.3 | 14.9± 9.3 |
| PRED-TMBB2<sup>4</sup> | 88.8±12.1 | 7.1±0.4 | - | - | 92.9±0.4 | 11.2±12.1 |
| PROFtmb | 91.9± 9.0 | 6.1±0.5 | - | - | 93.9±0.5 | 8.1± 9.0 |

\* Evaluation of the ability to distinguish between 57 beta barrel TMPs ( $\beta$ -TMP), 571 alpha helical TMPs ( $\alpha$ -TMP) and 5,654 globular, water-soluble non-TMP proteins in our data set. Recall and false positive rate (FPR) were averaged over the five independent cross-validation test sets; error margins given for the 95% confidence interval ( $1.96 \times$ standard error); **bold**: best values for each column; *italics*: differences statistically significant with over 95% confidence (only computed between best and 2<sup>nd</sup> best, or all methods ranked 1 and those ranked lower).
<sup>1</sup> Evaluation missing for one of 5,654 globular proteins.
<sup>2</sup> Evaluation missing for one of 571 $\alpha$ -TMPs and six of 5,654 globular proteins.
<sup>3</sup> Evaluation includes only 51 $\beta$ -TMPs, 552 $\alpha$ -TMPs, and 5,524 globular proteins due to runtime errors.
<sup>4</sup> The local PRED-TMBB2 version did not include the pre-filtering step of the web server. This caused a FPR for $\beta$ -TMP of almost 78%. Thus, we listed the statistics for the web server predictions, which did not include MSA input.

Many of the beta barrel TMPs that prediction methods missed had only two or four transmembrane beta strands (TMB). Such proteins cannot form a pore on their own, instead they have to form complexes with other proteins to function as TMPs, either by binding to other proteins or by forming multimers with additional copies of the same proteins by, e.g., trimerization. In fact, all four beta barrel TMPs missed by TMbed fell into this category. Thus, as all other methods, TMbed performed, on average, worse for beta barrel TMPs that cannot form pores alone. This appeared unsurprising, as the input to all methods were single proteins. For TMPs with TMHs, we also observed lower performance in the distinction between TMP/other for TMPs with a single TMH (recall: 93±3%) compared to those with multiple TMHs (recall: 99±1%). However, TMPs with single helices can function alone.

The embedding-based methods TMbed (introduced here using ProtT5 (34)) and DeepTMHMM (44) (based on ESM-1b (36)) performed at least on par with the SOTA using evolutionary information from MSA (Table 1). While this was already impressive, the real advantage was in the speed. For instance, our method, TMbed, predicted all 6,517 proteins in our data set in about 13 minutes (i.e., about eight sequences per second) on our server machine (SOM: Table S1); this runtime included generating the ProtT5 embeddings. The other embedding-based method, DeepTMHMM, needed about twice as long (23 minutes). Meanwhile, methods that search databases and generate MSAs usually take several seconds or minutes for a single protein sequence (70), or require significant amounts of computing resources (e.g., often more than 100 GB of memory) to achieve comparable runtimes (55).

### Excellent transmembrane segment prediction performance

TMbed reached the highest performance for transmembrane segments amongst all methods evaluated (Tables 2 & 3). With recall and precision values of 89±1% for TMHs, it significantly outperformed the second best and only other embedding-based method, DeepTMHMM, (80±2%, Table 2). TMbed essentially predicted 62% of all transmembrane helical (TMH) TMPs completely correctly (Q_ok_, i.e., all TMHs within ±5 residues of true annotation). DeepTMHMM reached second place with Q_ok_ of 46±4%. This difference between TMbed and DeepTMHMM was over twice that between DeepTMHMM and the two methods performing third-best by this measure, CCTOP (17, 18) and TOPCONS2 (29), which are based on evolutionary information.

**Table 2:**
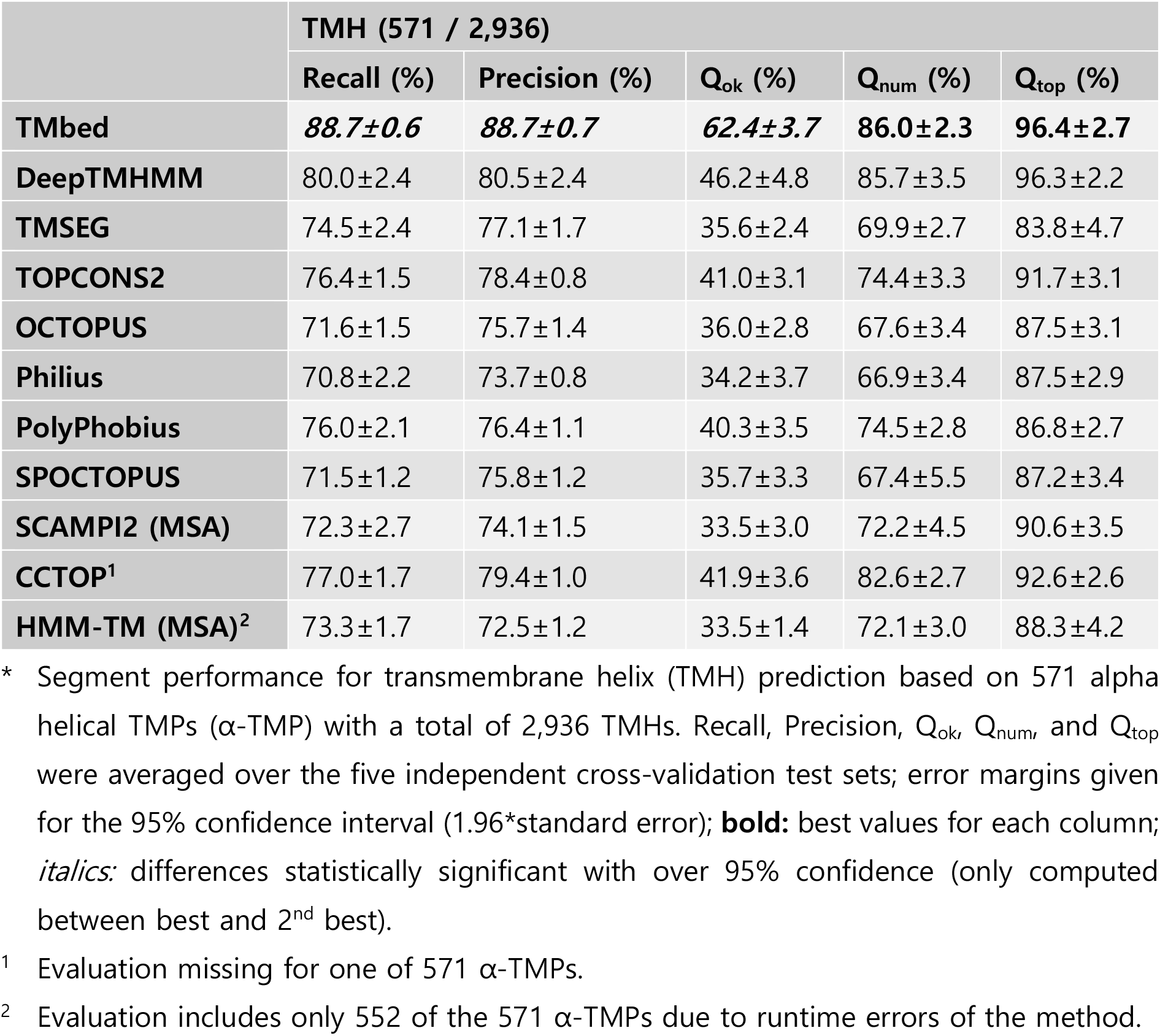
Per-segment performance for TMH (transmembrane helices). *.

**Table 3:**
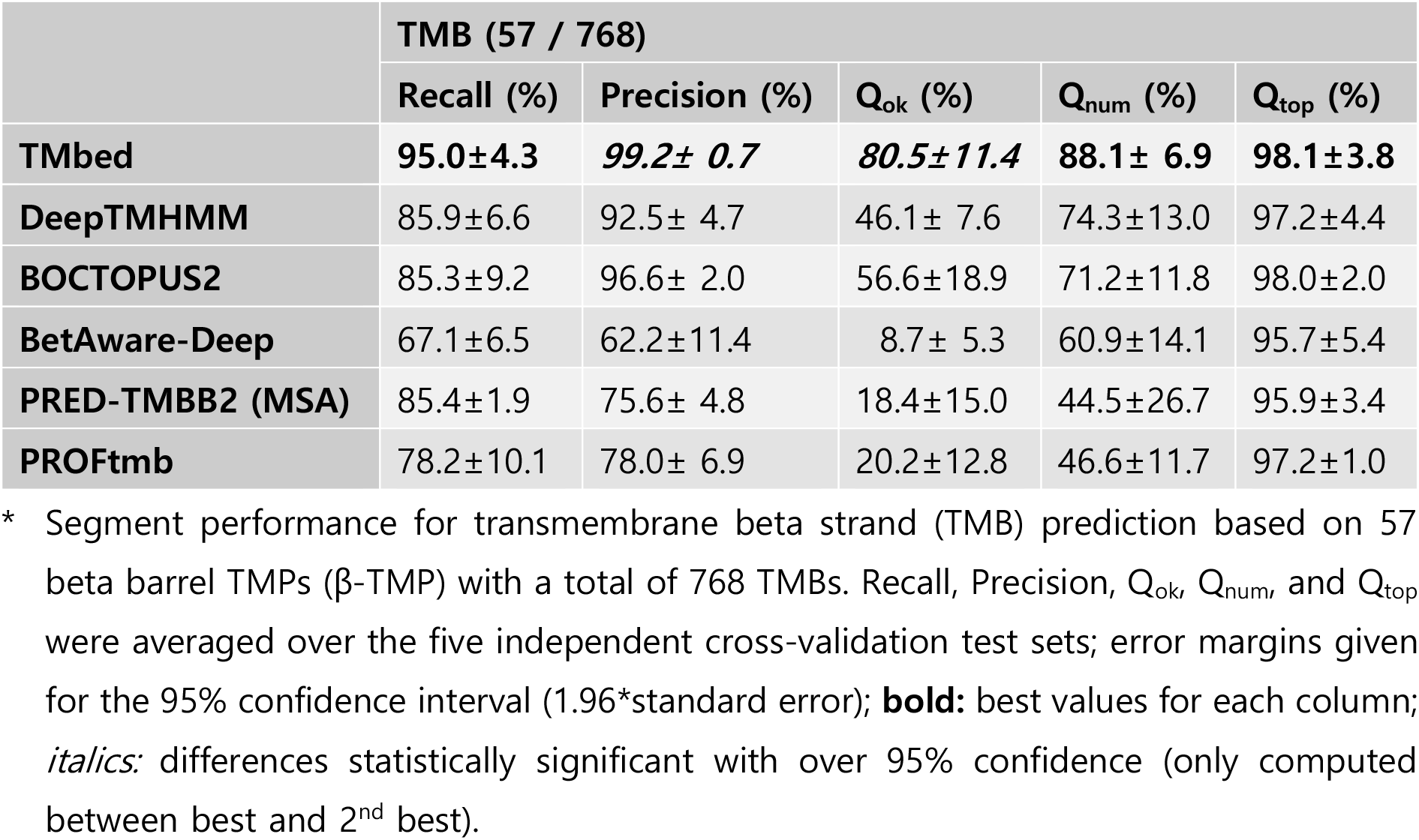
Per-segment performance for TMB (transmembrane beta strands). *.

The results were largely similar for beta barrel TMPs (TMBs) with TMbed achieving the top performance by all measures: reaching 95% recall and an almost perfect 99% precision. The most pronounced difference was a 23 percentage points lead in Q_ok_ with 80%, compared to BOCTOPUS2 (16) with 57% in second place. Overall, TMbed predicted the correct number of transmembrane segments in 86-88% of TMPs and correctly oriented 98% of TMBs and 96% of TMHs. For signal peptides, TMbed performed on par with SignalP 6.0, reaching 93% recall and 95% precision (SOM: Table S3). For this task, both methods appeared to be slightly outperformed by DeepTMHMM. However, none of those differences exceeded the 95% confidence interval, i.e., the numerically consistent differences were not statistically significant. On top, the signal peptide expert method SignalP 6.0 is the only of the three that distinguishes between different types of signal peptides. As for the overall per-protein distinction between TMP and non-TMP, the per-segment recall and precision also slightly correlated with the number of transmembrane segments, i.e., the more TMHs or TMBs in a protein the higher the performance (SOM: Table S4). Again, as for the TMP/non-TMP distinction, beta barrel TMPs with only two or four TMBs differed most to those with eight or more.

### Gaussian filter and Viterbi decoder improve segment performance

TMbed introduced a Gaussian filter smoothing over some local peaks in the prediction and a Viterbi decoder implicitly enforcing some “grammar-like” rules (Materials & Methods). We investigated the effect of these concepts by comparing the final TMbed architecture to three simpler alternatives: one variant used only the CNN, the other two variants combined the simple CNN with either the Gaussian filter or the Viterbi decoder, not both as TMbed. For the variants without the Gaussian filter, we retrained the CNN using the same hyperparameters but without the filter. Individually, both modules (filter and decoder) significantly improved precision and Q_ok_ for both TMH and TMB, while recall remained largely unaffected (SOM: Table S9). Clearly, either step already improved over just the CNN. However, which of the two was most important depended on the type of TMP: for TMH proteins Viterbi decoder mattered more, for TMB proteins the Gaussian filter. Both steps together performed best throughout without adding any significant overhead to the overall computational costs compared to the other components.

### Self-predictions reveal potential membrane proteins

We checked for potential overfitting of our model by predicting the complete data set with the final TMbed ensemble. This meant that four of the five models had seen each of those proteins during training. While the number of misclassified proteins went down, we found that there were still some false predictions, indicating that our models did not simply learn the training data by heart (SOM: Tables S7 & S8). In fact, upon closer inspection of the 11 false positive predictions (8 alpha helical and 3 beta barrel TMPs), those appear to be transmembrane proteins incorrectly classified as globular proteins in our data set due to missing annotations in UniProtKB/Swiss-Prot, rather than incorrect predictions. Two of them, P09489 and P40601, have automatic annotations for an autotransporter domain, which facilitates transport through the membrane. Further, we processed the predicted AlphaFold2 (11, 68) structures of all 11 proteins using the PPM (45) algorithm, which tries to embed 3D structures into a membrane bilayer. For eight of those, the predicted transmembrane segments correlated well with the predicted 3D structures and membrane boundaries (Fig. 1; SOM: Fig. S5). For the other three, the 3D structures and membrane boundaries still indicate transmembrane domains within those proteins, but the predicted transmembrane segments only cover parts of those domains (SOM: Fig. S5, last row). Together, these predictions provided convincing evidence for considering all eleven proteins as TMPs.

**Fig. 1:**
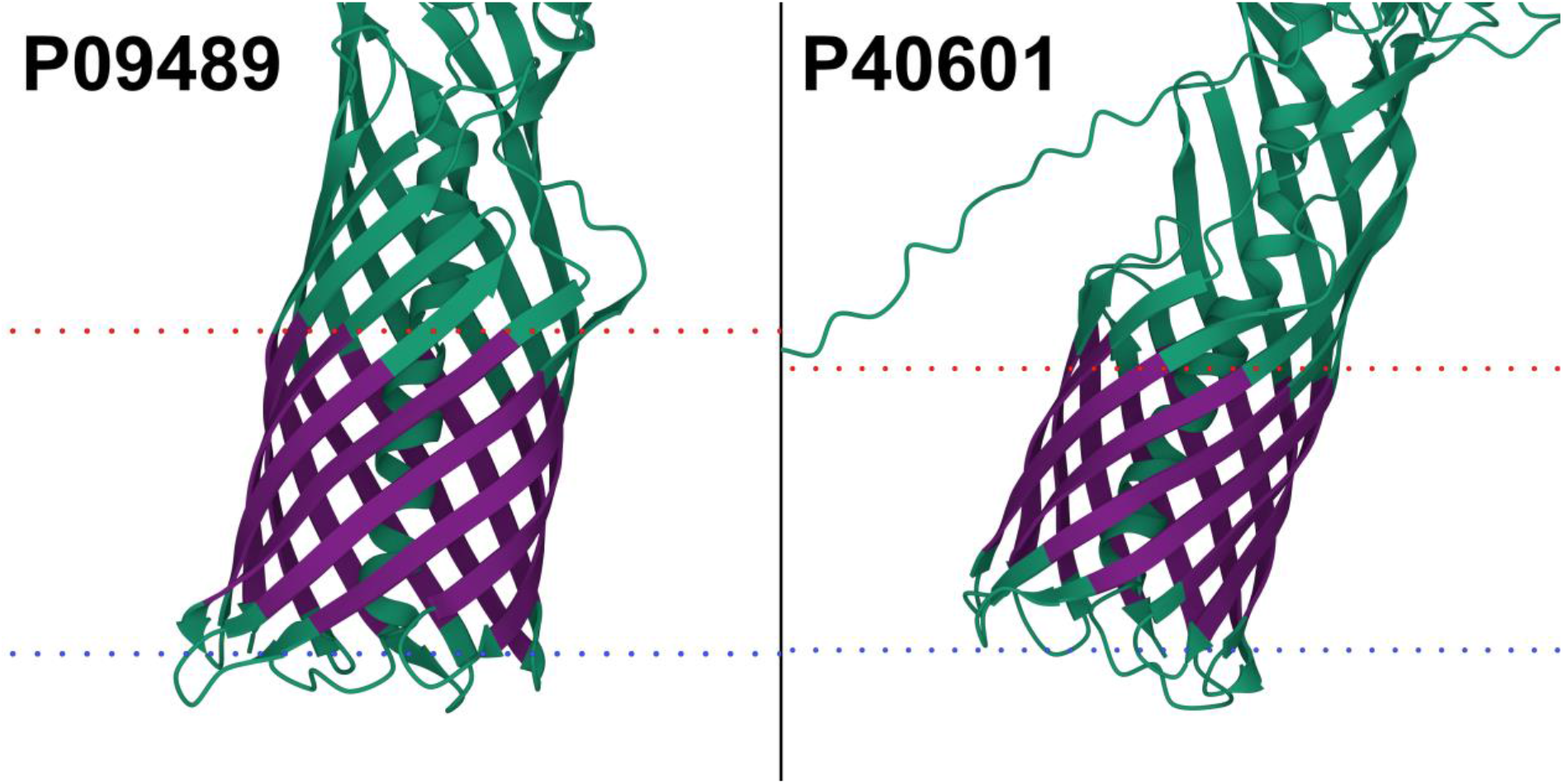
Potential transmembrane proteins in the globular data set. AlphaFold2 (11, 68) structure of extracellular serine protease (P09489) and Lipase 1 (P40601). Transmembrane segments (dark purple) predicted by TMbed correlate well with membrane boundaries (dotted lines: red=outside, blue=inside) predicted by the PPM (45) web server. Images created using Mol* Viewer (74). Though our data set lists them as globular proteins, the predicted structures indicate transmembrane domains, which align with segments predicted by our method. The predicted domains overlap with autotransporter domains detected by the UniProtKB (46) automatic annotation system. Transmembrane segment predictions were made with the final TMbed ensemble model.

### Predicting the human proteome in less than an hour

Given that our new method already outperformed the SOTA using evolutionary information from MSAs, the even more important advantage was speed. To estimate prediction throughput, we applied TMbed to all human proteins in 20,375 UniProtKB/Swiss-Prot (version: April 2022; excluding TITIN_HUMAN due to its extreme length of 34,350 residues). Overall, it took our server machine (SOM: Table S1) only 46 minutes to generate all embeddings and predictions (estimate for consumer-grade PC in the next section). TMbed identified 14 beta barrel TMPs and 4,953 alpha helical TMPs, matching previous estimates for alpha helical TMPs (1, 28). Two of the 14 TMBs appear to be false positives as TMbed predicted only a single TMB in each protein. The other 12 proteins are either part of the Gasdermin family (A to E), or associated with the mitochondrion, including three proteins for a voltage-dependent anion-selective channel and the TOM40 import receptor.

Further, we generated predictions for all proteins from UniProtKB/Swiss-Prot (version: May 2022), excluding sequences above 10,000 residues (20 proteins). Processing those 566,976 proteins took about 8.5 hours on our server machine. TMbed predicted 1,702 beta barrel TMPs and 77,296 alpha helical TMPs (predictions available via our GitHub repository).

### Hardware requirements

Our model needs about 2.5 GB of memory on the GPU when in 16-bit format. The additional memory needed during inference grows with the square of sequence length due to the attention mechanism of the transformer architecture. On our consumer-grade desktop PC (SOM: Table S1), this translated to a maximum sequence length of about 4,200 residues without maxing out the 12 GB of GPU memory. This barred 76 (0.4%) of the 20,376 human proteins from analysis on a personal consumer-hardware solution (NVIDIA GeForce RTX 3060). The prediction (including embedding generation) for 99.6% of the human proteome (20,376 proteins) took about 57 minutes on our desktop PC. While it is possible to run the model on a CPU, instead of on a GPU, we do not recommend this due to over 10-fold larger runtimes. More importantly, the current lack of support of 16-bit floating-point format on CPUs would imply doubling the memory footprint of the model and computations.

### Out-of-distribution performance

The two pLM-based methods DeepTMHMM (44) and TMbed appeared to reach similar performance according to the additional out-of-distribution data set (SOM: Tables S11 & S12). While DeepTMHMM reached higher scores for beta barrel proteins (Q_ok_ of 79±22% vs. 64±26%), these were not quite statistically significant. On the other hand, TMbed managed to outperform DeepTMHMM for alpha helical TMPs (Q_ok_ of 53±11% vs. 47±10%), though again without statistical significance. Furthermore, TMbed performed on par with the OPM baseline (SOM: Table S12), i.e., using the OPM annotations as predictions for the DeepTMHMM data set, implying that TMbed reached its theoretical performance limit on that data set. Surprisingly, TOPCONS2 and CCTOP both outperformed TMbed and DeepTMHMM with Q_ok_ of 65±10% and 64±10% (both not statistically significant), respectively.

Taking a closer look at the length distribution for the transmembrane segments in the TMbed and DeepTMHMM data set annotations and predictions (SOM: Fig. S6) revealed differences. First, while the TMB segments in both data sets averaged 9 residues in length, the DeepTMHMM distribution was slightly shifted toward shorter segments (left in Fig. S6A) but with a wider spread towards longer segments (right in Fig. S6A). Both of these features were mirrored in the distribution of predicted TMBs. In contrast, the TMH distributions for DeepTMHMM showed an unexpected peak for TMH with 21 residues (both in the annotations used to train DeepTMHMM and in the predictions). In fact, the peak for annotated TMHs at 21 was more than double the value of the two closest length-bins (TMH=20|22) combined. As the lipid bilayer remains largely invisible in X-ray structures, the exact begin and ends of TMHs may have some errors (28, 45, 49-51, 62). Thus, when plotting the distribution of TMH length, we expected some kind of normal distribution with a peak around 20-odd residues with more points for longer than for shorter TMHs (71). In stark contrast to this expectation, the distribution observed for the TMHs used to develop DeepTMHMM appeared to have been obtained through some very different protocol (Fig. S6B).

In contrast, the distributions for the annotations from OMP and the predictions from TMbed appeared to be more normally distributed with TMH lengths exhibiting a slight peak at 22 residues. The larger the AI model, the more it succeeds in reproducing features of the development set even when those might be based on less experimentally supported aspects. The DeepTMHMM model reproduced the dubious experimental distribution of TMHs exceedingly (Fig. S6B, e.g., orange line and bars around peak at 16). Although we do not know the origin of this bias in the DeepTMHMM data set, we have seen similar bias in some prediction methods and automated annotations in UniProtKB/Swiss-Prot. In fact, a quick investigation showed that for 80 of the 184 common alpha helical TMPs the DeepTMHMM annotations matched those found in UniProtKB but not the OPM annotation in our TMbed data set. Of those annotations, 66% (303 of 459) were 21-residues long TMHs, accounting for 73% of all such segments; the other 104 TMPs contained only 19% (114 of 593) TMHs of length 21. This led us to believe that the DeepTMHMM data set contained, in part, length-biased annotations found in UniProtKB. Other examples of methods with length biases include SCAMPI2 and TOPCONS2 that both predicted exclusively TMHs with 21 residues; OCTOPUS and SPOCTOPUS predicted only TMHs of length 15, 21, and 31 (with more than 90% of those being 21 residues). BOCTOPUS2 predicted only beta strands of length 8, 9, and 10, with about 80% of them being nine residues long.

Since TMHs are around 21 residues long, such bias is not necessarily relevant. However, it might point to why performance appears better against some data sets supported less by high-resolution experiments than by others.

### Performance on new membrane proteins

Although, the small data set size did not allow for statistically significant results (SOM: Table S13), TMbed performed numerically better than the other methods; in particular, BOCTOPUS2 failed to predict the only beta barrel TMP. While TMbed and DeepTMHMM both missed two of the 30 transmembrane beta strands, TMbed placed the remaining ones, on average, more accurately (recall: 93% vs 87%; precision: 100% vs. 93%). All methods performed worse for the alpha helical TMPs than on the other two benchmark data set, though with a sample size of only four proteins (25 TMHs total), we cannot be sure if this is an effect of testing on novel membrane proteins or simply by chance. Nevertheless, the transmembrane segments predicted by TMbed fit quite well to the membrane boundaries estimated by the PPM (63-65) algorithm (Fig. 2).

**Fig. 2:**
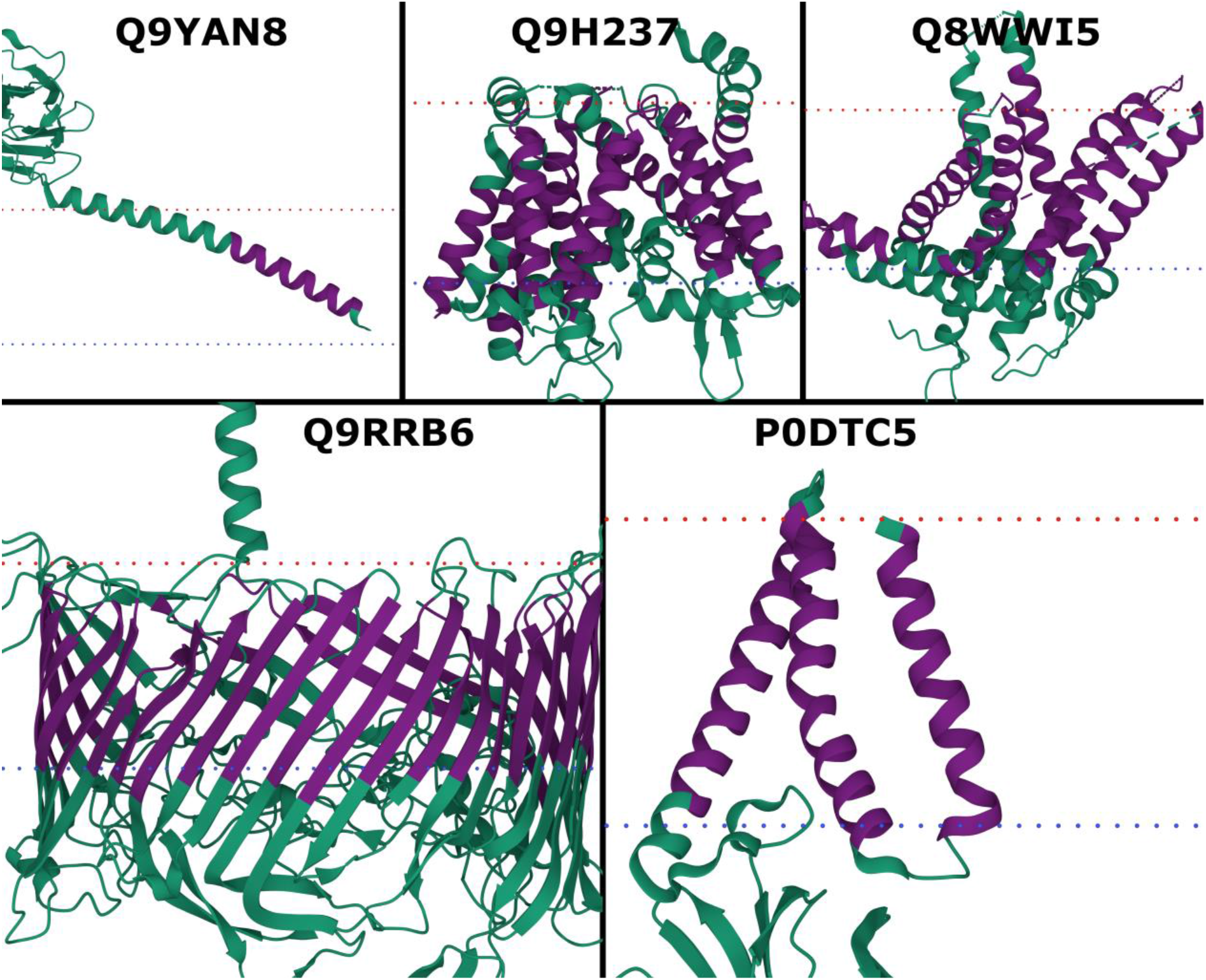
New membrane proteins. PDB structures for probable flagellin 1 (Q9YAN8; 7TXI (75)), protein-serine O-palmitoleoyltransferase porcupine (Q9H237; 7URD (76)), choline transporter-like protein 1 (Q8WWI5; 7WWB (77)), S-layer protein SlpA (Q9RRB6; 7ZGY (78)), and membrane protein (P0DTC5; 8CTK (79)). Transmembrane segments (dark purple) predicted by TMbed; membrane boundaries (dotted lines: red=outside, blue=inside) predicted by the PPM (45) web server. Images created using Mol* Viewer (74).

### No data leakage through pLM

pLMs such as ProtT5 (34) used by TMbed or ESM-1b (36) used by DeepTMHMM are pre-trained on billions of protein sequences. Typically, these include all protein sequences known today. In particular, they include all membrane and non-membrane proteins used in this study. In fact, assuming that the TMPs of known structure account for about 2-5% (72, 73) of all TMPs and that TMPs account for about 20-25% of all proteins, we assume pLMs have been trained on over 490 million TMPs that remain to be experimentally characterized. For the development of AI/ML solutions, it is crucial to establish that methods do not over-fit to existing data but that they will also work for new, unseen data. This implies that in the standard cross-validation process, it is important to not leak any data from development (training and validation used for hyperparameter optimization and model choice) to test set (used to assess performance). This implies the necessity for redundancy reduction. This also implies that the conditions for the test set are exactly the same as those that will be encountered in future predictions. For instance, if today’s experimental annotations were biased toward bacterial proteins, we might expect performance to be worse for eukaryotic proteins and *vice versa*.

Both TMbed introduced here and DeepTMHMM are based on the embeddings of pre-trained pLMs; both accomplish the TM-prediction through a subsequent step dubbed transfer learning, in which they use the pLM embeddings as input to train a new AI/ML model in supervised manner on some annotations about membrane segments. Could any data leak from the training of pLMs into the subsequent step of training the TM-prediction methods? Strictly speaking, if no experimental annotations are used, no annotations can leak: the pLMs used here never saw any annotation other than protein sequences.

Even when no annotations could have leaked because none were used for the pLM, should we still ascertain that the conditions for the test set and for the protein for which the method will be applied in the future are identical? We claim that we do not have to ascertain this. However, we cannot support any data for (nor against) this claim. To play devil’s advocate, let us assume we had to. The reality is that the vast majority of all predictions likely to be made over the next five years will be for proteins included in these pLMs. In other words, the conditions for future use-cases are exactly the same as those used in our assessment.

## Conclusions

TMbed predicts alpha helical (TMH) and beta barrel (TMB) transmembrane proteins (TMPs) with high accuracy (Table 1), performing at least on par or even better than state-of-the-art (SOTA) methods, which depend on evolutionary information from multiple sequence alignments (MSA; Tables 1-3). In contrast, TMbed exclusively inputs sequence embeddings from the protein language model (pLM) ProtT5. Our novel method shines, in particular, through its low false positive rate (FPR; Table 1), incorrectly predicting fewer than 1% of globular proteins to be TMPs. TMbed also numerically outperformed all other tested methods in terms of correctly predicting transmembrane segments (on average, 9 out of 10 segments were correct; Tables 2 & 3). Despite its top performance, the even more significant advantage of TMbed is speed: the high throughput rate of the ProtT5 (34) encoder enables predictions for entire proteomes within an hour, given a suitable GPU (SOM: Table S1). On top, the method runs on consumer-grade GPUs as found in more recent gaming and desktop PCs. Thus, TMbed can be used as a proteome-scale filtering step to scan for transmembrane proteins. Validating the predicted segments with AlphaFold2 (11, 68) structures and the PPM (45) method could be combined into a fast pipeline to discover new membrane proteins, as we have demonstrated with a few proteins. Finally, we provide predictions for 566,976 proteins from UniProtKB/Swiss-Prot (version: May 2022) via our GitHub repository.

## Declarations

### Ethics approval and consent to participate

Not applicable.

### Consent for publication

Not applicable.

### Availability of data and materials

Our code, method, and data sets are freely available in the GitHub repository, https://github.com/BernhoferM/TMbed.

### Authors’ contributions

MB collected the data sets, developed and evaluated the TMbed model, and took the lead in writing the manuscript. BR supervised and guided the work, and co-wrote the manuscript. All authors read and approved the final manuscript.

## Acknowledgements

Thanks to Tim Karl and Inga Weise for their help with technical and administrative issues; to Tobias Olenyi, Michael Heinzinger, and Christian Dallago for thoughtful discussions, help with ProtT5, and help with the manuscript; to Konstantinos Tsirigos and Ioannis Tamposis for their support with setting up HMM-TM and PRED-TMBB2; to Pier Luigi Martelli for providing us with BetAware-Deep predictions. Thanks to all who deposit their experimental data in public databases, and to those who maintain them. Last but not least, we thank the reviewers for their constructive criticism, which helped to improve our manuscript.

## Abbreviations used

CI: confidence interval
CNN: convolutional neural network
MSA: multiple sequence alignment
OPM: orientations of proteins in membranes database
PDB: protein data bank
PDBTM: protein data bank of transmembrane proteins
pLM: protein language model
SIFTS: structure integration with function, taxonomy and sequence
SOM: supplementary online material
SOTA: state-of-the-art
SP: signal peptide
TMB: transmembrane beta strand
TMH: transmembrane helix
TMP: transmembrane protein

