## Supplementary Online Material for "TMbed – Transmembrane proteins predicted through Language Model embeddings"

#### Table of Contents for Supporting Online Material

### Short description of Supporting Online Material

Here, we provide short explanations for the different performance metrics used during evaluation (Note S1, Fig. S4); how we created the results for several other prediction methods (Note S2); we briefly explain the idea behind using a simple Viterbi decoder and its limitations (Note S3, Fig. S2); and we provide illustrative sketches of our model architecture (Fig. S1) and nested cross-validation process (Fig. S3).

We list the hardware specifications of the machines used during the project (Table S1) and the optimal hyperparameters for each of the final models (Table S2). Further, we provide performance statistics for signal peptides (Table S3) and protein groups based on their number of transmembrane segments (Table S4). We also list statistics for the individual models and cross-validation splits (Tables S5 & S6), confusion matrices for the cross-validation and final models (Table S7 & S8), the effect of the Gaussian filter and Viterbi decoder on segment performance (Table S9), and estimate the expected number of mistakes made in a hypothetical proteome (Table S10). Additionally, we show a few more “false positives” that might actually be transmembrane proteins (Fig. S5). Finally, we provide performance and annotation statistics for an out-of-distribution data set gathered from DeepTMHMM (Tables S11 & S12, Fig. S6) and a CASP-like data set of novel membrane proteins (Table S13).

### Material

#### Note S1: Performance metrics explained.

We evaluated our and other methods using several standard and non-standard performance metrics, listed below. Statistics referring to a specific type of segment (i.e., transmembrane beta strands or helices, and signal peptides) are calculated using only the corresponding subset of proteins. For example, the *precision* and $Q_{ok}$ values for transmembrane helices take only the 571 alpha helical transmembrane proteins (TMPs) into account, ignoring any false positive predictions made in beta barrel TMPs or globular proteins.

*Recall,* also called Sensitivity, is the percentage of positive samples (proteins or segments) that have been correctly predicted as such. For example, TMbed correctly identified 557 of the 571 alpha helical TMPs, i.e., a *recall* of about 98%.

$$Recall = \frac{Number\ of\ correct\ positive\ predictions}{Number\ of\ positive\ samples} * 100\% \qquad \textit{(Eq. S1)}$$

*Precision* reflects the percentage of positive predictions that are actually correct. Since only 557 of the 584 alpha helical TMPs predicted by TMbed are correct (the other 27 are globular proteins), it has a *precision* of about 95%.

$$Precision = \frac{Number\ of\ correct\ positive\ predictions}{Number\ of\ positive\ predictions} * 100\% \qquad \textit{(Eq. S2)}$$

*False Positive Rate (FPR)* gives the percentage of negatives samples incorrectly predicted as positive ones. For example, 27 out of 5711 globular and beta barrel TMPs incorrectly predicted as alpha helical TMPs correspond to a *FPR* of about 0.5%.

$$FPR = \frac{Number\ of\ incorrect\ positive\ predictions}{Number\ of\ negative\ samples} * 100\% \qquad \textit{(Eq. S3)}$$

$Q_{ok}$ is the percentage of proteins for which all predicted segments of a given type are correct, i.e., segment *recall* and *precision* are both 100% for those proteins. A $Q_{ok}$ of 79% for beta barrel TMPs corresponds to 45 of 57 proteins that do not have any false positive or false negative predictions of transmembrane beta strands.

$$Q_{ok} = \frac{Number\ of\ proteins\ with\ all\ segment\ of\ type\ T\ correct}{Number\ of\ proteins\ with\ segments\ of\ type\ T} * 100\% \qquad \textit{(Eq. S3)}$$

$Q_{num}$ shows the percentage of proteins that have the correct number of predicted segments of a given type, regardless of their exact position. TMbed predicted the correct number of transmembrane beta strands in 50 of 57 beta barrel TMPs, i.e., it has a $Q_{num}$ of about 88%.

$$Q_{num} = \frac{Number\ of\ proteins\ with\ correct\ number\ of\ segments\ of\ type\ T}{Number\ of\ proteins\ with\ segments\ of\ type\ T} * 100\% \qquad \textit{(Eq. S4)}$$

$Q_{top}$ gives the percentage of correctly predicted segments of a given type that also have the correct inside/outside orientation, i.e., its endpoints are on the correct sides of the membrane. For example, TMbed correctly predicts 730 of 768 transmembrane beta strands (*recall* of about 95%). Of those 730 segments, 714 also have the correct inside/outside orientation, i.e., the $Q_{top}$ value is about 98%. We consider only the first residue on each side of a segment to determine its orientation.

$$Q_{top} = \frac{Number\ of\ correct\ segments\ of\ type\ T\ with\ correct\ orientation}{Number\ of\ correct\ segments\ of\ type\ T} * 100\% \qquad \textit{(Eq. S5)}$$

We estimate the error margin of our performance values with the 95% confidence interval (CI), i.e., 1.96 times the standard error (SE) based on the sample standard deviation (SD):

$$CI = 1.96 * SE; \quad SE = \frac{SD}{\sqrt{N}}; \quad SD = \sqrt{\frac{1}{N-1} * \sum_{i=1}^{N} (x_i - \bar{x})^2}, \qquad \textit{(Eq. S6-S8)}$$

where $N$ is the number of measurements $x_i$ performed and $\bar{x}$ is the mean over those. In our case, $N$ usually refers to the five cross-validation iterations.

---

**Note S2: Other prediction methods.**

In order to put the performance of TMbed into context, we made predictions for the proteins in our data set using several other methods.

DeepTMHMM (1) uses ESM-1b (2) embeddings to predict alpha helical and beta barrel transmembrane proteins (TMP). We generated all predictions using a local installation as described on their web server homepage (https://dtu.biolib.com/DeepTMHMM).

TOPCONS2 (3), OCTOPUS (4), Philius (5), PolyPhobius (6), and SPOCTOPUS (7) all predict alpha helical TMPs. TOPCONS2, Philius, PolyPhobius, and SPOCTOPUS additionally predict signal peptides. With the exception of Philius, all other five methods use evolutionary information in the form of BLAST profiles or MSAs as additional input to the protein sequence. As TOPCONS2 is a consensus prediction method combining all of the above methods, we got all predictions from its web server (https://topcons.net). Unfortunately, the web server rejected one of the globular proteins, P05790, due to its high GA content (incorrectly thought to be a DNA sequence).

CCTOP (8, 9) is another consensus prediction method for alpha helical TMPs. It combines a total of 10 prediction methods and topology constraint determined by a homology lookup. We used their web server (https://cctop.ttk.hu/) to generate predictions for our data sets. Due to sequence length restrictions (up to 5,000 residues) we are missing predictions for one alpha helical TMP and six globular proteins.

SCAMPI2 (10) is an improved version of the older SCAMPI (11) method employed as part of TOPCONS2. We downloaded the software from its GitHub repository[1] and used UniRef90 as the BLAST search database to generate the alignments needed for the MSA version of SCAMPI2.

HMM-TM (12) and PRED-TMBB2 (13) are methods predicting alpha helical and beta barrel TMPs, respectively. We computed predictions for our data set using their respective web servers (http://www.compgen.org/tools). For both methods, we used the most recent improvements employing hidden neural networks (14). As the online services only allow batch submissions for the single sequence versions, i.e., without the use of MSAs as input, we also installed local versions of their methods (15) and

[1] https://github.com/ElofssonLab/scampi2

ran them offline. However, the offline version of PRED-TMBB2 does not include the protein filtering using pHMMs that the web server employs, significantly increasing its false positive rate. Unfortunately, the local MSA versions failed for some of the proteins and we were unable to fix the issue. Thus, we are missing predictions for 155 proteins (6 beta barrel TMPs, 19 alpha helical TMPs, 130 globular proteins) by HMM-TM (MSA) and 26 proteins (2 alpha helical TMPs, 24 globular proteins) by PRED-TMBB2 (MSA).

The authors of BetAware-Deep (16) kindly provided us with predictions for our data set as the web server only allows for submission of a single sequence at a time. Their method combines sequence profiles with several machine learning architectures (LSTM, CRF) to predict beta barrel TMPs.

We installed and ran an offline version of BOCTOPUS2 (17) to predict beta barrel TMPs in our data set. We generated the sequence profiles and results according to the descriptions on their GitHub repository[2].

For TMSEG (18) and PROFtmb (19) we used the predictions generated by our PredictProtein (20) pipeline. TMSEG and PROFtmb predict alpha helical and beta barrel TMPs, respectively, both using BLAST profiles as additional input.

We used the SignalP 6.0 (21) web server[3] to generate additional signal peptide predictions. We chose the “slow” model mode to get accurate predictions. Just like TMbed, SignalP 6.0 uses a protein language model to generate embeddings (22).

**Note S3: Viterbi decoder.**

We use an untrained Viterbi decoder to translate the class probability distributions generated by our models into actual class labels for each residue in a sequence. The decoder scores state transitions according to the class probabilities predicted by the CNN model, trying to find the path with the highest sum of probabilities. We apply a score penalty of $-100$ to transitions not intended by our defined grammar (Fig. S2), effectively preventing the decoder from considering those transitions.

The main purpose of the decoder is to enforce a small set of rules:

1) Signal peptides may only start at the N-terminus of a sequence.
2) Signal peptides and transmembrane segments must be at least five residues long.
3) The inside/outside orientation of non-membrane parts must change after every transmembrane segment.

We explicitly model the transition from IN to OUT state to allow for sequence parts that pass the membrane boundaries without actually being in contact with the membrane. For example, this includes parts of beta barrel TMPs that pass through the pore formed by their own beta barrel structure (Manuscript: Fig. 1).

2 https://github.com/ElofssonLab/boctopus2

3 https://services.healthtech.dtu.dk/service.php?SignalP-6.0

A downside to this is that the model is free to change the IN/OUT state to accommodate transmembrane segments even if the majority of the non-membrane residues on both sides of the segment are on the same side of the membrane, circumventing rule 3.

For example: `...iiiiiHHHHHHHHHHHoiiii...`

However, disallowing the direct transitions between IN and OUT would prevent correctly modelling sequences where such transitions do happen and can encourage the model to split transmembrane segments to insert a single non-membrane residue, thereby accommodating the same orientation on both sides of the original segment.

For example: `...iiiiiHHHHHHoHHHHHiiii...`

Given those two alternatives, we decided to allow for direct transitions.

**Table S1: Hardware specifications.**

| | CPU | RAM | GPU | VRAM | Storage |
|---|---|---|---|---|---|
| **Desktop Machine** | Intel Core i5-2500K | 24GB DDR3 | NVIDIA GeForce RTX 3060 | 12GB GDDR6 | SSD |
| **Server Machine** | Intel Xeon Gold 6248 | 400GB DDR4 ECC | NVIDIA Quadro RTX 8000 | 48GB GDDR6 | SSD |

* List of most relevant hardware components in the two machines used during method development. We used the server machine to create sequence embeddings, while training and testing our new method on the Desktop machine.

**Table S2: Final hyperparameters.**

| | Learning rate | Weight decay |
|---|---|---|
| **Model 0** | 0.005 | 0.01 |
| **Model 1** | 0.010 | 0.10 |
| **Model 2** | 0.005 | 0.10 |
| **Model 3** | 0.010 | 0.10 |
| **Model 4** | 0.010 | 0.10 |

* Optimal learning rate and weight decay values selected for each model during nested cross-validation. Numbers in the model name indicate the test set; for example, model 0 was trained and optimized on sets 1-4, and evaluated on set 0.

**Table S3: Signal peptide performance.**

| | Proteins (661 vs. 4993) | | Segments (661) | |
|---|---|---|---|---|
| | Recall (%) | FPR (%) | Recall (%) | Precision (%) |
| **TMbed** | 98.8±0.8 | **0.1±0.1** | 93.3±1.4 | 94.5±1.3 |
| **SignalP 6.0** | 97.0±1.0 | 0.2±0.1 | 92.6±1.7 | 95.5±1.7 |
| **DeepTMHMM** | **99.2±0.5** | 0.2±0.2 | **95.3±1.8** | **96.0±1.7** |
| **TMSEG** | 95.0±1.5 | 5.3±0.4 | 77.4±3.7 | 81.5±3.7 |
| **TOPCONS2**[1] | 93.9±1.7 | 2.3±0.5 | 81.8±3.0 | 87.1±2.3 |
| **Philius**[1] | 93.3±1.6 | 5.9±0.6 | 84.5±1.8 | 90.6±1.7 |
| **PolyPhobius**[1] | 92.4±1.7 | 1.9±0.3 | 84.1±2.7 | 91.0±1.8 |
| **SPOCTOPUS**[1] | 93.5±1.8 | 3.6±0.5 | 85.3±2.7 | 91.2±1.5 |

* Protein and segment performance for signal peptide (SP) prediction based on 661 globular proteins with SPs and 4993 globular proteins without SPs. Performance values were averaged over the five independent cross-validation test sets; error margins given for the 95% confidence interval (1.96*standard error); **bold:** best values for each column; *italics:* differences statistically significant with over 95% confidence (only computed between best and 2nd best).

[1] Evaluation includes only 660 of the 661 globular proteins with SPs due to one sequence being rejected by the prediction web server.

**Table S4: Performance relative to number of transmembrane segments.**

| | TMB | | TMH | | |
|---|---|---|---|---|---|
| | 2, 4 (13) | 8+ (44) | 1 (164) | 2-5 (175) | 6+ (232) |
| **$Q_{ok}$ (%)** | 65.0±36.7 | 85.6±8.6 | 78.1±3.1 | 61.0±5.5 | 52.7±3.2 |
| **Recall (%)** | 65.0±36.7 | 96.6±4.4 | 79.3±4.5 | 84.1±1.9 | 90.6±0.7 |
| **Precision (%)** | 75.0±38.0 | 99.5±0.6 | 83.4±2.7 | 83.8±2.3 | 90.3±0.8 |

* TMbed segment performance for transmembrane beta strand (TMB) and helix (TMH) prediction based on 57 beta barrel and 571 alpha helical TMPs. TMPs are subdivided into groups based on their number of transmembrane segments: a) 2 or 4 TMBs, b) 8 or more TMBs, c) a single TMH, d) 2-5 TMHs, and e) 6 or more TMHs. The numbers in parenthesis indicate the number of proteins within that group. Performance values were averaged over the five independent cross-validation test sets; error margins given for the 95% confidence interval (1.96*standard error).

**Table S5: Cross-Validation protein performance.**

| | β-TMP | | α-TMP | | Globular | |
|---|---|---|---|---|---|---|
| | Recall (%) | FPR (%) | Recall (%) | FPR (%) | Recall (%) | FPR (%) |
| **Model 0** | 85.7 | 0.1 | 97.5 | 0.2 | 99.7 | 3.7 |
| **Model 1** | 100.0 | 0.0 | 97.3 | 0.7 | 99.3 | 2.5 |
| **Model 2** | 83.3 | 0.0 | 96.5 | 0.7 | 99.3 | 4.8 |
| **Model 3** | 100.0 | 0.2 | 98.2 | 0.5 | 99.3 | 1.6 |
| **Model 4** | 100.0 | 0.0 | 98.3 | 0.3 | 99.7 | 1.6 |
| **Average ±1.96SE** | *93.8±7.5* | *0.1±0.1* | *97.5±0.7* | *0.5±0.2* | *99.5±0.2* | *2.8±1.2* |

* Protein prediction performance for each TMbed model on the corresponding independent test set. Numbers in the model name indicate the test set; for example, model 0 was trained on sets 1-4 and evaluated on set 0. Last row lists the average over all five sets and 1.96 times the sample standard error, i.e., the 95% confidence interval.

**Table S6: Cross-Validation TM segment performance.**

| | TMB | | | TMH | | |
|---|---|---|---|---|---|---|
| | $Q_{ok}$ (%) | Recall (%) | Precision (%) | $Q_{ok}$ (%) | Recall (%) | Precision (%) |
| **Model 0** | 64.3 | 95.6 | 98.5 | 56.7 | 88.7 | 88.4 |
| **Model 1** | 83.3 | 87.1 | 98.4 | 68.2 | 89.6 | 89.3 |
| **Model 2** | 75.0 | 94.9 | 99.2 | 61.1 | 88.4 | 89.3 |
| **Model 3** | 80.0 | 97.6 | 100.0 | 62.8 | 87.8 | 87.5 |
| **Model 4** | 100.0 | 100.0 | 100.0 | 63.5 | 88.9 | 89.1 |
| **Average ±1.96SE** | *80.5±11.4* | *95.0±3.4* | *99.2±0.7* | *62.4±3.7* | *88.7±0.6* | *88.7±0.7* |

* Segment prediction performance for each TMbed model on the corresponding independent test set. Numbers in the model name indicate the test set; for example, model 0 was trained on sets 1-4 and evaluated on set 0. Last row lists the average over all five sets and 1.96 times the sample standard error, i.e., the 95% confidence interval.

##### Table S7: Cross-Validation confusion matrix.

| | | Predicted Label | | | |
|---|---|---|---|---|---|
| | | **β-TMP** | **α-TMP** | **SP** | **No SP** |
| **True Label** | **β-TMP** | 53 | 0 | 3 | 1 |
| | **α-TMP** | 0 | 557 | 4 | 10 |
| | **SP** | 3 | 9 | 643 | 6 |
| | **No SP** | 0 | 18 | 6 | 4969 |

* Aggregated confusion matrix for the five TMbed models on the corresponding independent test sets. Protein categories are beta barrel TMPs (β-TMP), alpha helical TMPs (α-TMP), globular proteins with signal peptides (SP) and globular proteins without signal peptides (No SP).

##### Table S8: Ensemble confusion matrix.

| | | Predicted Label | | | |
|---|---|---|---|---|---|
| | | **β-TMP** | **α-TMP** | **SP** | **No SP** |
| **True Label** | **β-TMP** | 56 | 0 | 1 | 0 |
| | **α-TMP** | 0 | 567 | 0 | 4 |
| | **SP** | 3 | 1 | 656 | 1 |
| | **No SP** | 0 | 7 | 4 | 4982 |

* Confusion matrix for the final TMbed ensemble on the complete data set, i.e., for every sequence there are four models that have seen it during training and one model that has not. Protein categories are beta barrel TMPs (β-TMP), alpha helical TMPs (α-TMP), globular proteins with signal peptides (SP) and globular proteins without signal peptides (No SP).

**Table S9: Effect of Gaussian filter and Viterbi decoder.**

| | TMB | | | TMH | | |
|---|---|---|---|---|---|---|
| | $Q_{ok}$ (%) | Recall (%) | Precision (%) | $Q_{ok}$ (%) | Recall (%) | Precision (%) |
| **TMbed** | 80.5±11.4 | 95.0±4.3 | 99.2±0.7 | 62.4±3.7 | 88.7±0.6 | 88.7±0.7 |
| **Viterbi** | 77.7±15.9 | 95.4±4.0 | 99.1±1.1 | 61.6±3.5 | 88.9±0.8 | 88.7±0.8 |
| **Gaussian** | 79.1±13.5 | 95.3±3.8 | 98.2±1.6 | 57.6±4.3 | 86.1±0.7 | 84.7±1.2 |
| **CNN** | 45.5±13.9 | 94.7±4.0 | 92.0±6.2 | 50.0±4.7 | 87.4±1.0 | 79.6±2.5 |

* Comparison between the CNN model, models combining the CNN with either the Gaussian filter or the Viterbi decoder, and the final TMbed model combining all three components. Segment performances for transmembrane beta strand (TMB) and helix (TMH) prediction are based on 57 beta barrel and 571 alpha helical TMPs. Performance values were averaged over the five independent cross-validation test sets; error margins given for the 95% confidence interval (1.96*standard error).

##### Table S10: Expected misclassifications.

| | Misclassifications | |
|---|---|---|
| | β-TMP | α-TMP |
| **TMbed** | 22 | 193 |
| **DeepTMHMM** | 53 (+31) | 278 (+85) |
| **TMSEG** | | 521 (+328) |
| **TOPCONS2** | | 683 (+490) |
| **OCTOPUS** | | 1666 (+1473) |
| **Philius** | | 766 (+573) |
| **PolyPhobius** | | 933 (+740) |
| **SPOCTOPUS** | | 2701 (+2508) |
| **SCAMPI2 (MSA)** | | 1135 (+942) |
| **CCTOP** | | 744 (+551) |
| **HMM-TM (MSA)** | | 3352 (+3159) |
| **BOCTOPUS2** | 880 (+858) | |
| **BetAware-Deep** | 984 (+962) | |
| **PRED-TMBB2** | 1573 (+1551) | |
| **PROFtmb** | 1049 (+1027) | |

* Number of expected misclassified proteins in a hypothetical proteome with 20,000 proteins. Misclassifications are the sum of false positive and false negative predictions for the specific tasks of predicting proteins with transmembrane alpha helices (α-TMP) or beta strands (β-TMP). The number in parentheses is the difference to our method, TMbed. Expected prediction error rates for the methods are based on their confusion matrices on our data set of 6282 proteins. The hypothetical proteome contains 5000 (25%) alpha helical and 200 (1%) beta barrel transmembrane proteins.

**Table S11: Out-of-distribution protein performance.**

| | β-TMP | | α-TMP | | Globular | |
|---|---|---|---|---|---|---|
| | **Recall (%)** | **FPR (%)** | **Recall (%)** | **FPR (%)** | **Recall (%)** | **FPR (%)** |
| **TMbed** | **100.0±0.0** | 0.6±0.6 | 95.5±4.5 | ***1.2±0.9*** | 98.1±1.2 | 3.9±3.9 |
| **DeepTMHMM** | **100.0±0.0** | **0.5±0.5** | 95.4±4.4 | ***1.2±0.9*** | **98.2±1.1** | 3.9±3.8 |
| **TOPCONS2** | - | - | 98.8±2.4 | 6.8±2.0 | 93.2±2.0 | 1.2±2.4 |
| **CCTOP**[1] | - | - | **98.9±2.2** | 5.5±1.9 | 94.5±1.9 | 1.1±2.2 |
| **BOCTOPUS2** | **100.0±0.0** | 5.9±1.8 | - | - | 94.1±1.8 | **0.0±0.0** |

* Evaluation of the ability to distinguish between 14 beta barrel TMPs (β-TMP), 86 alpha helical TMPs (α-TMP) and 567 globular, water-soluble non-TMP proteins from the DeepTMHMM data set; all proteins are non-redundant with respect to the TMbed data sets. Recall and false positive rate (FPR) were averaged over 1000 bootstrap iterations (random sampling with replacement); error margins given for the 95% confidence interval (1.96*standard deviation); **bold:** best values for each column; *italics:* differences statistically significant with over 95% confidence (only computed between best and 2nd best, or all methods ranked 1 and those ranked lower).

1 Evaluation missing for one of 85 α-TMPs.

#### Table S12: Out-of-distribution segment performance.

| | TMB | | | TMH | | |
|---|---|---|---|---|---|---|
| | $Q_{ok}$ (%) | Recall (%) | Precision (%) | $Q_{ok}$ (%) | Recall (%) | Precision (%) |
| **TMbed** | 64.2±25.7 | 93.3±5.3 | 93.3±5.3 | 52.6±10.8 | 82.8±4.7 | 83.8±4.8 |
| **DeepTMHMM** | **78.8±22.4** | **98.2±1.8** | **98.2±1.8** | 46.8±10.4 | 79.8±5.5 | 80.0±5.6 |
| **TOPCONS2** | - | - | - | **65.0±10.0** | ***88.9±3.7*** | **89.4±3.7** |
| **CCTOP[1]** | - | - | - | 63.5±10.0 | **88.9±4.0** | 89.0±3.9 |
| **BOCTOPUS2** | 43.0±26.1 | 91.0±5.3 | 92.0±4.9 | - | - | - |
| **OPM[2]** | 61.1±14.3 | 92.0±6.4 | 91.6±6.4 | 52.7±7.2 | 83.9±2.9 | 83.3±3.0 |

* Segment performance for transmembrane beta strand (TMB) and helix (TMH) prediction based on 14 beta barrel and 86 alpha helical TMPs from the DeepTMHMM data set; all proteins are non-redundant with respect to the TMbed data sets. $Q_{ok}$, recall and precision were averaged over 1000 bootstrap iterations (random sampling with replacement); error margins given for the 95% confidence interval (1.96*standard deviation); **bold:** best values for each column; *italics:* differences statistically significant with over 95% confidence (only computed between best and 2nd best, or all methods ranked 1 and those ranked lower; ignores the OPM baseline).

1 Evaluation missing for one of 85 α-TMPs.

2 OPM represents the baseline for how much the DeepTMHMM data set annotations agree with our annotations collected from the OPM database, i.e. we are using the OPM annotations as predictions for the DeepTMHMM data set. The performance statistics were evaluated for a set of 44 beta barrel and 184 alpha helical TMPs common to both data sets (TMbed and DeepTMHMM).

**Table S13: Segment performance on new membrane proteins.**

| | TMB | | | TMH | | |
|---|---|---|---|---|---|---|
| | $Q_{ok}$ (%) | Recall (%) | Precision (%) | $Q_{ok}$ (%) | Recall (%) | Precision (%) |
| **TMbed** | **0.0** | **93.3** | **100.0** | **25.0** | **60.0** | **62.5** |
| **DeepTMHMM** | **0.0** | 86.7 | 92.9 | **25.0** | 40.0 | 43.5 |
| **TOPCONS2** | - | - | - | **25.0** | 52.0 | 48.1 |
| **CCTOP** | - | - | - | **25.0** | 48.0 | 50.0 |
| **BOCTOPUS2** | **0.0** | 0.0 | 0.0 | - | - | - |

* Segment performance for transmembrane beta strand (TMB) and helix (TMH) prediction based on one beta barrel and four alpha helical TMPs; all proteins are non-redundant with respect to the TMbed data sets and the DeepTMHMM data set. Error margins omitted due to extremely small data set size; none of the performance differences are statistically significant; **bold:** best values for each column.

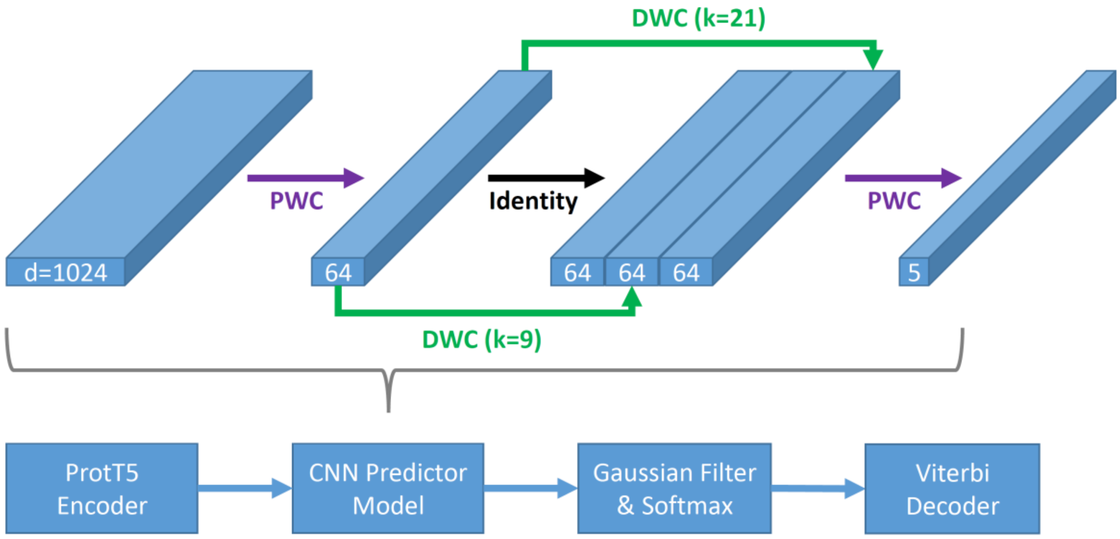

**Figure S1: TMbed model architecture.**

The TMbed model consists of four parts: a) The ProtT5 encoder converts the input sequence into per-residue embeddings with 1024 dimensions for each residue in the sequence; b) a convolutional neural network (CNN) predicts class scores based on those embeddings; c) a Gaussian filter smooths the class scores and converts them into class probabilities via the softmax function; d) a Viterbi decoder assigns class labels to each of the residues in the sequence. The CNN consists of four layers: two pointwise convolutions (PWC) and two depthwise convolutions (DWC; kernel sizes of 9 and 21). The output of the first PWC and both DWCs also passes through layer normalization and a ReLU activation function.

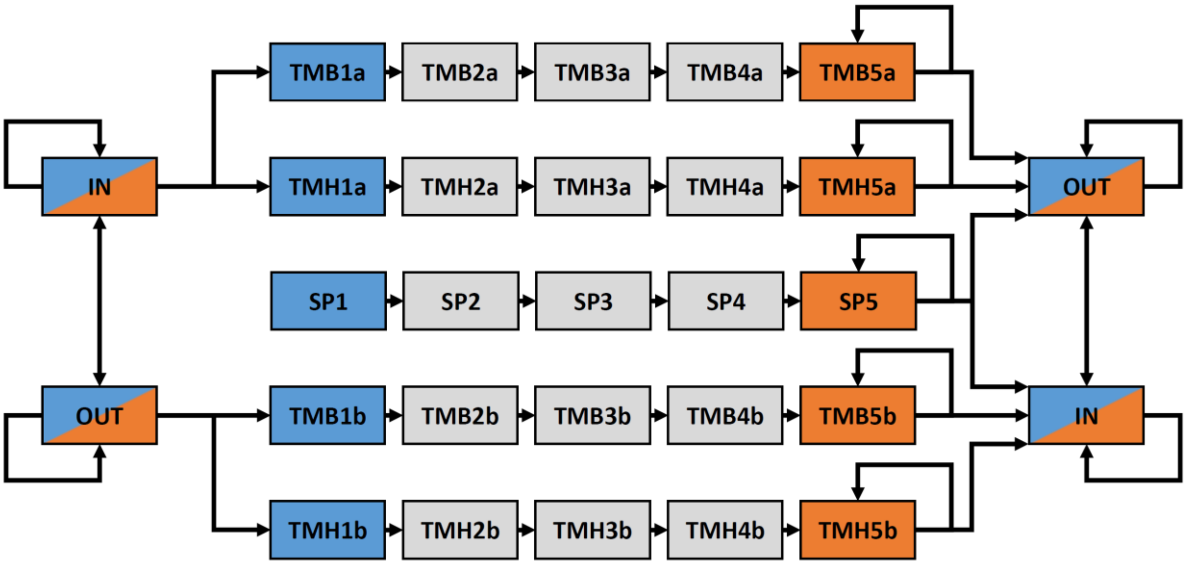

**Figure S2: Viterbi decoder state transitions.**

Transitions encoded in the Viterbi decoder to go from one state to another state. We split transmembrane beta strands (TMB), helices (TMH), and signal peptides (SP) into sub-states to enforce minimum segment lengths of five residues. A decoded sequence must start with one of the blue states and may only end with one of the orange states. The IN and OUT states on both sides represent the same two internal states and are only duplicated to simplify the graph.

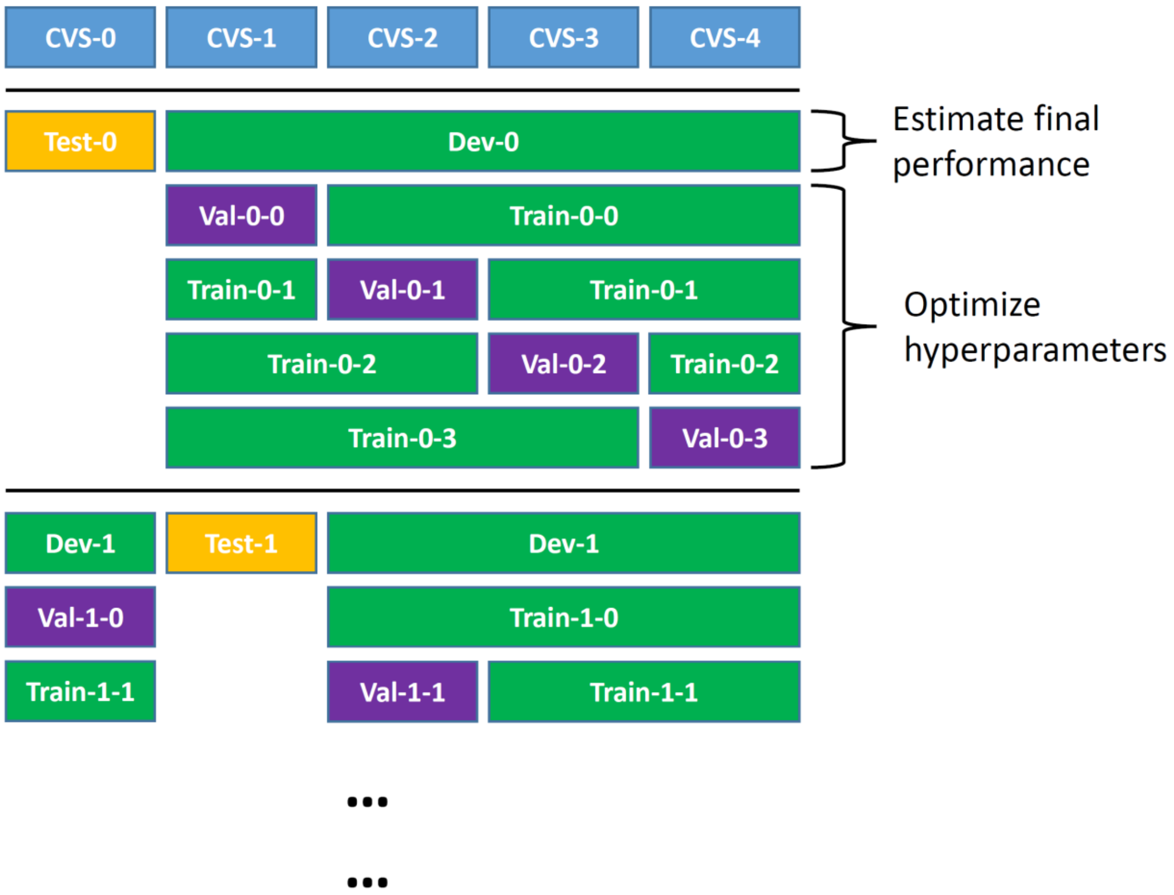

**Figure S3: Nested Cross-Validation.**

For the nested cross-validation process, we split the data set into five cross-validation splits (CVS). During each of the outer five iterations, we used one split as the test set to estimate the models final performance and the other four to develop the model. We further divided those four splits into training set and validation set. We then trained the model on the training set and evaluated the performance on the validation set, repeating the process for each hyperparameter combination. We repeated this process three more times, each time using a different split for the validation set. We chose the best set of hyperparameters based on the average performance on all four validation sets, trained the model using those parameters on the development set, and evaluated its performance on the test set. We repeated this overall process four more times, each time choosing a different CVS for the test set, until each CVS had been used for testing once. This process yielded five trained models, which we used for the final TMbed ensemble.

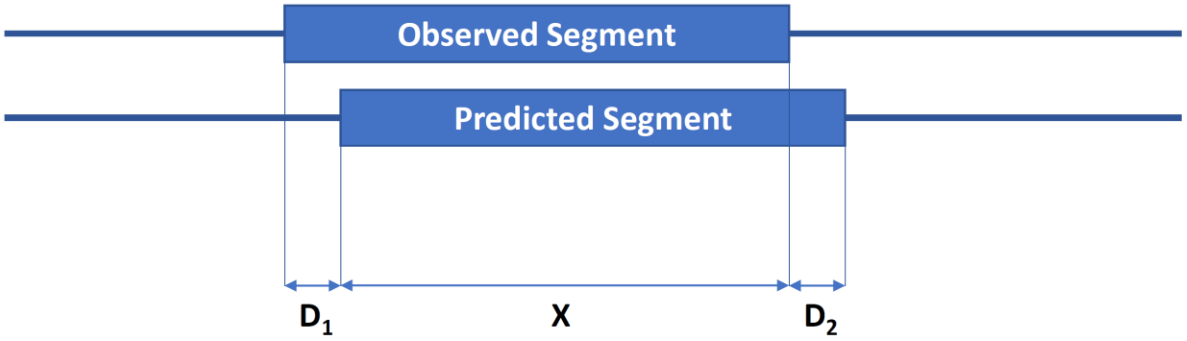

**Figure S4: Segment validation criteria.**

Illustration of the two criteria for a predicted segment to be correct: 1) start and end positions must not deviate by more than five residues, i.e., $\max(D_1, D_2) \leq 5$, and 2) the intersection (overlap) between the observed and predicted segment must be at least half of their union, i.e., $\frac{X}{X+D_1+D_2} \geq 0.5$.

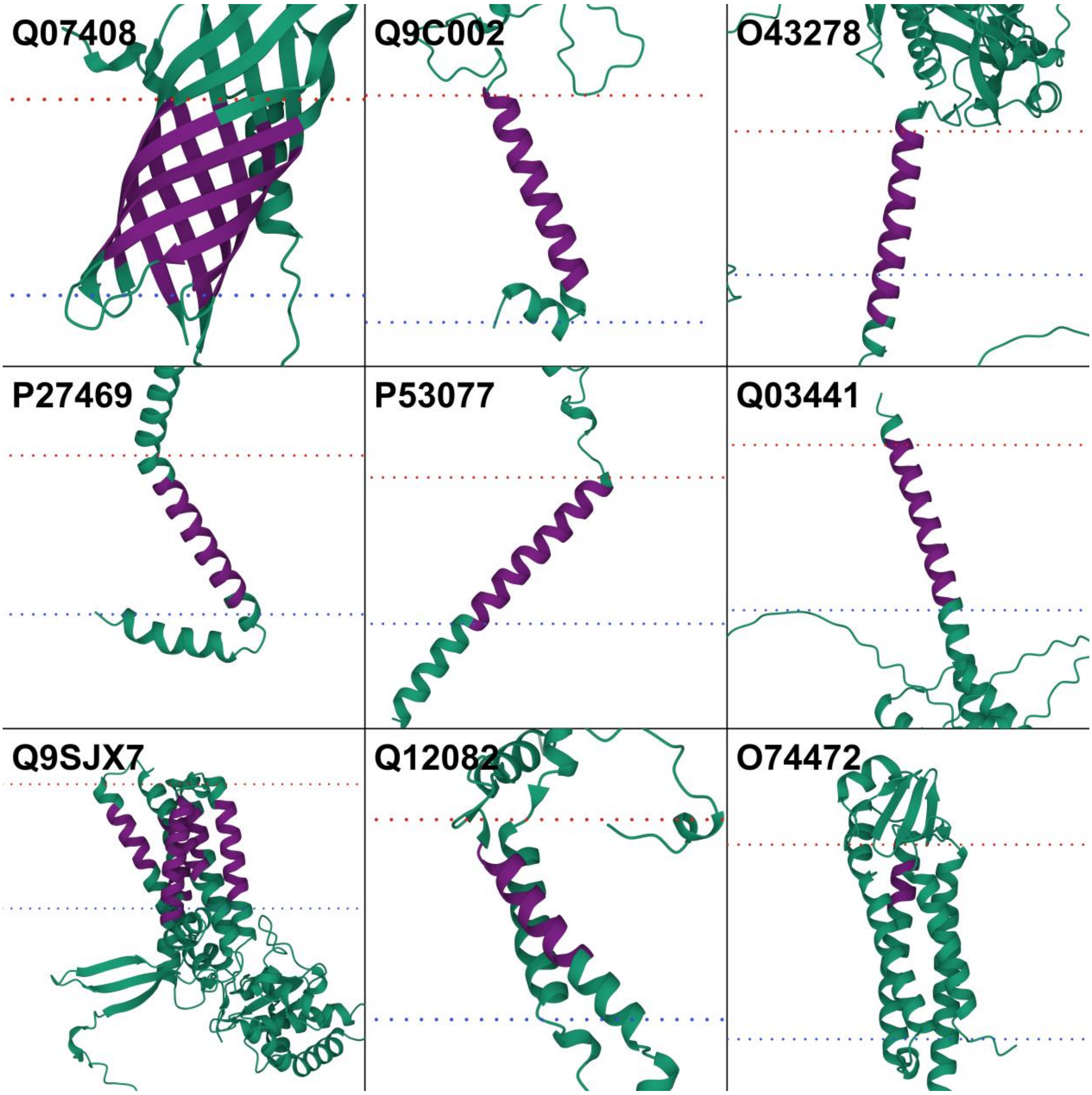

**Figure S5: More potential transmembrane proteins in the globular data set.**

AlphaFold2 (23, 24) structures of nine proteins from the globular data set: major surface antigen 4 (Q07408), normal mucosa of esophagus-specific gene 1 protein (Q9C002), Kunitz-type protease inhibitor 1 (O43278), G0/G1 switch protein 2 (P27469), maintenance of telomere capping protein 3 (P53077), sporulation protein RMD1 (Q03441), protein root UVB sensitive 2 (Q9SJX7), uncharacterized protein YDL157C (Q12082), and meiotically up-regulated gene 33 protein (O74472). For most proteins, transmembrane segments (dark purple) predicted by TMbed correlate well with membrane boundaries (dotted lines: red=outside, blue=inside) predicted by the PPM (25) web server. Images created using Mol* Viewer (26). Though our data set lists them as globular proteins, the predicted structures indicate transmembrane domains, which align with segments predicted by our method. Predictions were made with the final TMbed ensemble model.

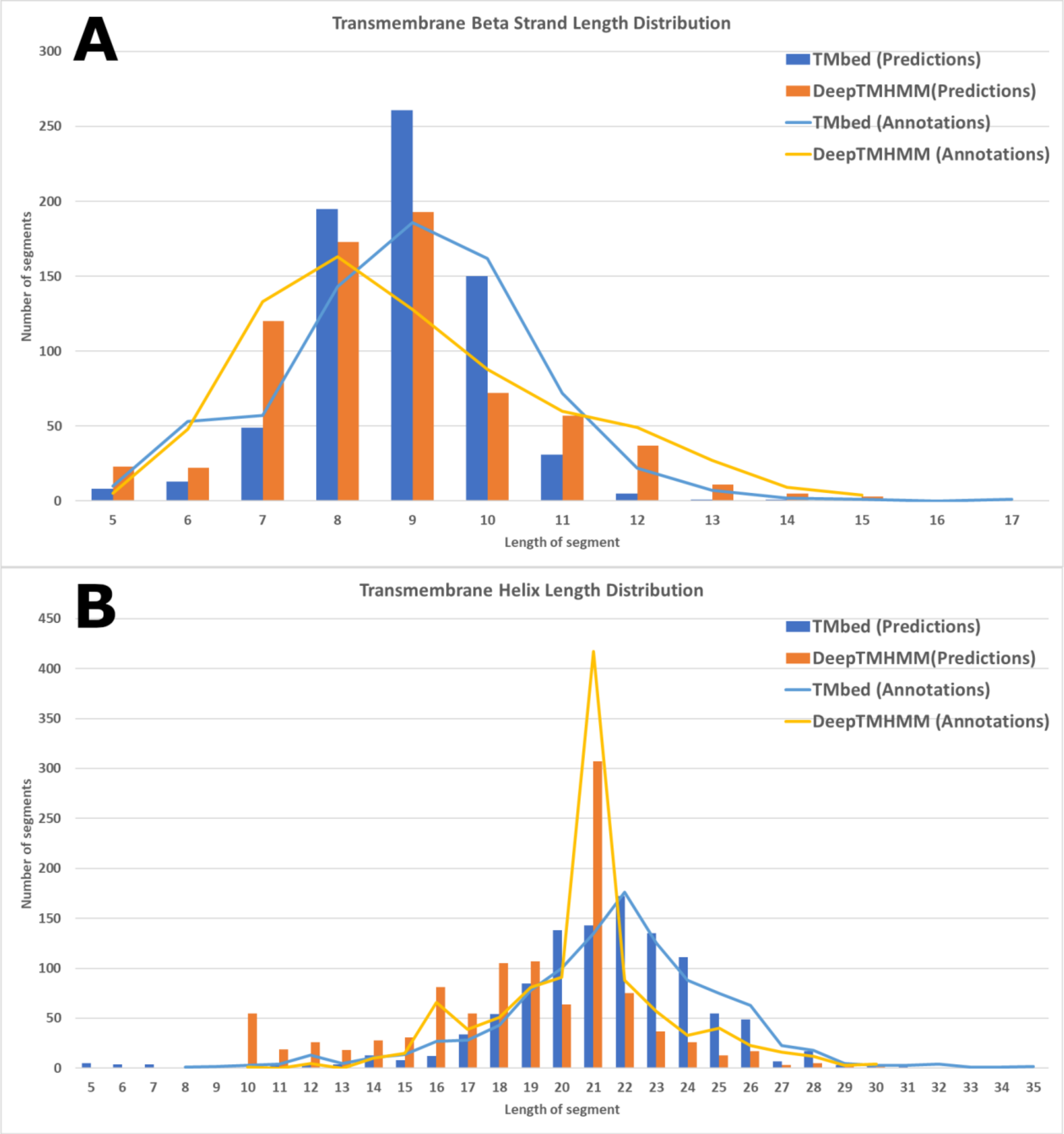

**Figure S6: Out-of-distribution segment length statistics.**

Transmembrane segments length distributions for 44 beta barrel (**A**) and 184 alpha helical (**B**) transmembrane proteins common to both the TMbed and DeepTMHMM data sets. **Lines:** statistics for the annotated segments in each data set; **Bars:** statistics for the segments predicted by each method during its individual cross-validation. Panel B is cropped to the right, missing two annotated segments (L: 40, 44) and four predicted segments (L: 37, 38, 40, 43) for TMbed.
